# Transcriptome analysis of fasting caecotrophy on hepatic lipid metabolism in New Zealand rabbits

**DOI:** 10.1101/493957

**Authors:** Yadong Wang, Huifen Xu, Guirong Sun, Mingming Xue, Shuaijie Sun, Tao Huang, Jianshe Zhou, Juan J Loor, Ming Li

**Author notes:** Author to whom correspondence should be addressed; **Ming Li**: E-Mail; **Juan J Loor**. These authors contributed equally to this paper (Yadong Wang, Huifen Xu).

## Abstract

**Background:** Rabbit produce two kinds of feces: hard and soft feces, and they have a preference for consuming the latter. Although this habit of rabbits has been reported for many years, little is known on whether this behavior will impact growth performance and metabolism. The RNA-Seq technology is an effective means of analyzing transcript groups to clarify molecular mechanisms. The aim of the present study was to investigate the effects of fasting caecotrophy on growth performance and lipid metabolism in rabbits.

**Results:** Our results indicated that, compared with the control group, the final body weight, weight gain, liver weight, specific growth rate and feed conversion ratio were all decreased in the experimental group (*P*<0.05). Oil red staining of the liver tissue indicated that fasting caecotrophy resulted in decrease of lipid droplet accumulation. RNA sequencing (RNA-seq) analysis revealed a total of 301.2 million raw reads approximately 45.06 Gb of high-quality clean data. The data were mapped to the rabbit genome (http://www.ensembl.org/Oryctolagus_cuniculus). After a five step filtering process, 14964 genes were identified, including 444 differentially expressed genes (*P*<0.05, foldchange≥1). Especially for remarkable changes of genes related to lipid metabolism, RT-PCR further validated the remarkable decrease of these genes in fasting caecotrophy group, including CYP7A1, PPARG, ABCA1, ABCB1, ABCG1, GPAM, SREBP, etc. KEGG annotation of the differentially expressed genes indicated that the main pathways affected were retinol metabolism, pentose and glucuronide interactions, starch and sucrose metabolism, fatty acid degradation, steroid hormone biosynthesis.

**Conclusion:** In conclusion, the present study revealed that banning caecotrophy reduced growth rate and altered lipid metabolism, our results laid instructive basis for rabbit feeding and production. These data also provides a reference for studying the effects of soft feces on other small herbivores.

## Introduction

Many small herbivores have a natural instinct of caecotrophy[1]. Because of the small body shape and their digestive tract volume is limited, the average residence time of food in the digestive tract is relatively short [2, 3]. In order to meet their nutrition needs, small herbivores need to obtain adequate high-quality food[4]. Small herbivores mainly rely on low-quality, high-fiber plant stems and leaves as food sources, with cellulose from symbiotic microorganisms in the hind-gut aiding in digestion[5, 6]. Because microbial fermentation takes longer time than the average residence time of food in the digestive tract, increasing digestibility by ingesting incompletely-digested nutrients is an important nutritional strategy for small herbivores [7–9].

There are two types of feces excreted by rabbits: hard feces (nutrient-poor) and soft feces (consist of protein, vitamins, and inorganic salts) [10–12]. Soft feces also contain a large number of microorganisms, which are important for microbial fermentation in herbivores. Thus, we speculate that rabbits might build their intestinal microbial flora through their caecotrophy behavior[13–15]. Although modern feeding and management techniques are designed to fully meet the nutritional needs for growth, rabbits still maintain the habit of eating soft feces[16, 17]. One study comparing high and low body weight Rex rabbits, the high-weight group had more abundant bacterial groups than the low-weight group, with bacterial groups in soft feces being more abundant than hard feces[18–20]. Intestinal flora and their host have a mutually beneficial symbiosis, with the host nutrition and metabolism, immune system development, disease resistance and other physiological functions playing an important role[21, 22].

Liver plays an important role in overall metabolism of lipids, including its function in supporting digestion, absorption, decomposition, synthesis and transportation of lipids[23, 24]. Bile acids produced by the liver are conversion products of cholesterol metabolism. These compounds are important for emulsifying dietary lipids and promote the digestion and absorption of lipids[25]. By generating ATP for its own utilization, the liver is also the main site for oxidative decomposition of fatty acids and production of ketone bodies[26–28]. In monogastrics, liver is the primary site for fatty acids oxidation and ketone bodies production[29, 30]. The liver-synthesized cholesterol accounts for more than 80% of the total cholesterol produced by the body, and is the main source of plasma cholesterol [31].

Transcriptome sequencing is the study of the gene specific expression in the same cell at different times or under different treatment conditions[32]. Because of its advantages for the high accuracy, high throughput, high sensitivity and low cost, it has been widely used in researches[33]. By using transcriptome sequencing we can better understand the expression of all genes in the liver, and then make a comprehensive analysis of genes that affect lipid metabolism, providing quality assurance for our data [34].

One study showed that caecrophagy affected the body weight of Rex rabbits. Whether caecotrophy affects the growth performance and lipid metabolism of the New Zealand white rabbits has not been reported. In order to investigate the effects of fasting caecotrophy on lipid metabolism and its possible mechanism, we performed feeding experiment and subject their liver tissues to transcriptomic sequencing, aimed to clarify the effects of fasting caecotrophy on rabbits growth rate and lipid metabolism, thus provide experimental evidence for rabbit breeding and production.

## Materials and methods

### 2.1 Animals and experimental design

#### 2.1.1 Ethic statement

The present study was designed and performed according to the guidelines of institutional Animals Care and Use Committee College of Animal Husbandry and Veterinary Medicine of Henan Agricultural University.

#### 2.1.2 Animals

Twelve, 28-day-old female weaned New Zealand white rabbits from a breeding center (western suburbs breeding filed of rabbit, Zhengzhou, China) were raised in the animal experimental center of Henan Agriculture University. The experimental room and cage was disinfected before raising the rabbit. Thirty-six rabbits with similar body weight (1.14±0.12kg) were randomly divided into two groups, with18 replicates in each group. Group A was a control group and was fed normally; group B was experimental group wearing Elizabeth circle to prevent caecotrophy. (Fig.1) The temperature was controlled at around 23±1 °C and rabbits had free access to food and water. Specifically, they were fed with the same pelleted diet at the breeding farm, and the diet was replaced with the experimental diet gradually.

**Fig 1.**
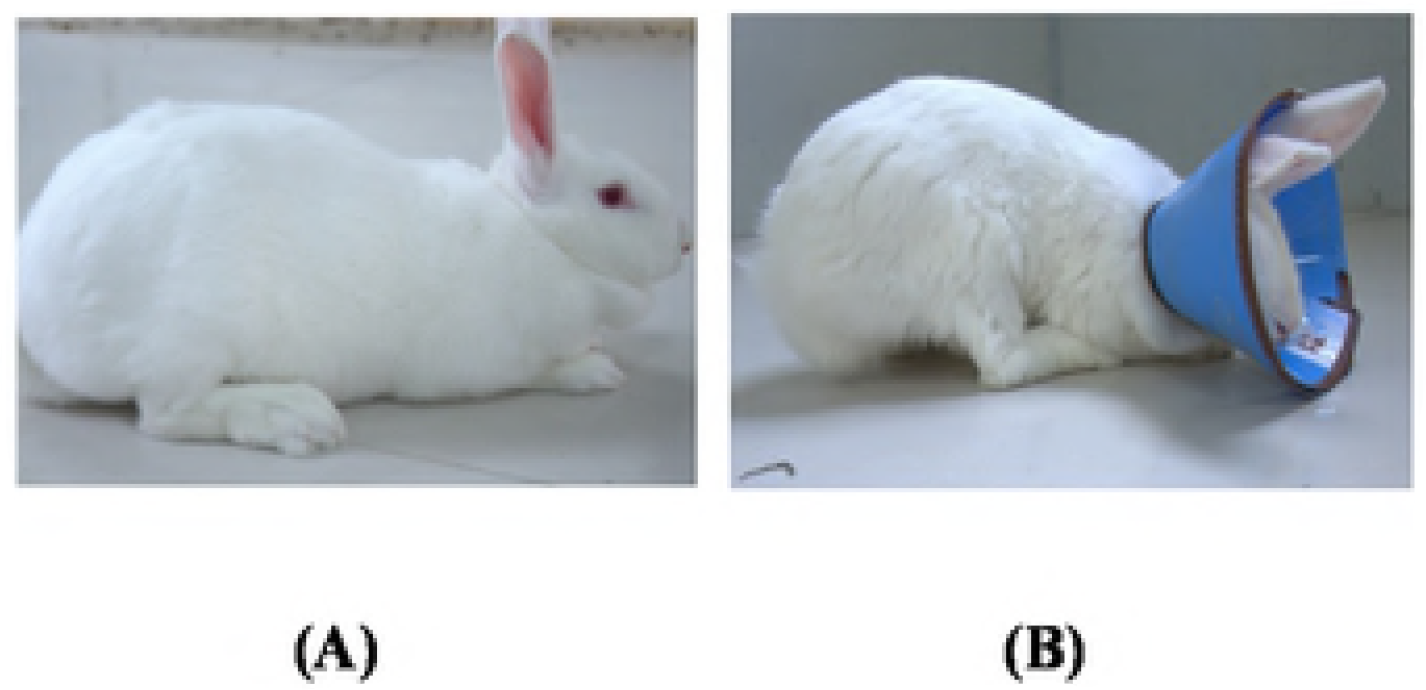
Rabbits of the two groups. (A) Rabbits of control group. (B) Rabbits of the experimental group. Rabbits of control group is feed normal, rabbits of the experimental group which wear a Elizabeth circle to prevent eat soft feces.

#### 2.1.3 Sample collection

After a 90-day long experiment period, twelve rabbits were anesthetized and sacrificed for sample extraction. Blood samples were collected in a tube with EDTA for clinical biochemistry measurements with standard protocol (JianCheng Bioengineering Institute, Nanjing, China). Liver, spleen, kidney, lung, heart, muscle and adipose were collected and weighted, and were put into liquid nitrogen immediately until analysis. Body weight and feed intake was recorded every week. These data were determined to analyze changes in various growth indices, liver and kidney index.

#### 2.1.4 Histology

Liver tissue from twelve rabbits were fixed with paraformaldehyde, embedded in paraffin, and then cut into 5 μm sections. These sections were subjected for oil red O staining to measure lipid droplet accumulation, lipid droplets were observed under Nikon microscope (Nikon Eclipse Ti-SR) and then quantified by Image-Pro Plus 6.0 software (Media Cybernetics, USA).

#### 2.1.5 Blood biochemical parameters

Blood biochemical parameters were analyzed according to manufacturer’s protocol (JianCheng Bioengineering Institute, Nanjing, China). A Super microplate reader (Bio-tek ELx808™) were used to measure high density lipoprotein cholesterol (HDL), low density lipoprotein cholesterol (LDL), total protein (TP) and albumin (ALB). The Malate Dehydrogenase (MDH) of blood and liver tissues were measured on an ultraviolet spectrophotometer according to manufacturer’s protocol (JianCheng Bioengineering Institute, Nanjing, China).

### 2.2 Transcriptomic analysis

Total RNA was extracted from the liver with Trizol reagent(Invitrogen,USA), according to the mannufacture’s instruction, then re-suspended in 50 μg RNase-free water and stored at -80 °C. The quality and concentration of the total RNA was quantified with a NanoDrop 2000 spectrophotometer (Thermo scientific). RNA integrity was accessed using the Nano 6000 LabChip kit(Agilent,Technologies, CA, USA) with RIN number >8.6.

#### 2.2.1 cDNA library preparation and Illumina sequencing

The high-quality total RNA of the rabbit livers (n = 3 per group) were send for the preparation of RNA-seq libraries. All the libraries were sequenced by the Illumina HiSeq™ 2500 platform (Biomarker technologies Corporation, Beijing, China).

#### 2.2.2 De novo assembly of sequencing reads

Raw Data (raw reads) of the Fastq format were first processed through in-house per scripts. Clean data were obtained by removing reads containing adapter, reads containing poly-N and low quality reads from raw data. There were 45.06 Gb clean data in all. Reads in which more than 91.53% of base have a Q value ≥ 30 were filtered out with the Fastq filter software (Biomarker Technologies)(S1 Fig). The clean reads were mapped to the reference genome of the rabbit using the TopHat2 software, which have an alignment efficiency>80%, (S2 Fig). Redundant sequences were eliminated, and the longest transcripts were recognized as unigenes, which were grouped together for the final assembly and subsequent annotation.

#### 2.2.3 Gene function Annotation

Gene function was annotated based on the following databases:Nr(NCBI non-redundant protein sequence); Nt(NCBI non-redundant nucleotide sequence); Pfam(Protein family); KOG/COG(Clusters of Orthologous Groups of proteins); Swiss-prot(A manually annotated and reviewed sequence database); KO(KEGG Ortholog database); GO(gene Ontology). Quantification of gene expression levels were estimated by fragments per kilobase of million fragments mapped. Differential expression analysis of two groups was performed using the DESeq R package. DESeq provide statistical routines for determining differential expression in digital gene expression data using a model based on the negative binomial distribution. The resulting P value was using the Benjamini and Hochberg’s approach for controlling the false discovery rate. Here, only unique reads with an absolute value of log^2^ ratio ≥ 1 and FDR significance score < 0.01 were used for the subsequent analysis.

Based on the TopHat2 alignment of Reads and reference sequences of each sample, the Cufflinks software was used for splicing and expression quantification. By comparison with known genome annotation files, a new transcriptome region, the new gene, is identified. The new gene was aligned with each database by software BLAST to obtain annotation information of new genes.

### 2.3 qRT-PCR

Quantitative real-time PCR (qRT-PCR) was used to measure the expression of 10 differential expressed genes to validate our illumina sequencing data. Total RNA was extracted from liver of rabbits in control and experimental groups (with six replicate in each group). Primers designed based on the assembled transcriptome sequence using the primer 6 software (primer biosoft International) and are listed in the S3 Table. The first strand cDNA was synthesized from 1 μg of RNA by using PrimeScript™ RT reagent Kit with gDNA Eraser (TaKaRa,Japan). All cDNA products were diluted to 200 ng/μL. The 20 μL qRT-PCR reaction mixture consisted of 2 μL template cDNA, 0.4μL of each primer, 10 μL of TB Green™ Premix Ex Taq™ (2X), 0.4 μL ROX and 6.8 μL of nuclease-free water. PCR amplification was performed in a 96-well optical plate at 95°C for 2 min, followed by 40 cycles of 95°C for 30s, 60°C for 30s, and a final extension at 72°C for 2 min. qRT-PCR was performed using the StepOne plus real-Time PCR system (Applied Biosystems) and 2^-ΔΔCT^ method was used tocalculate the relative expression level of each genes. β-actin gene was used as the reference gene for normalization.

### 2.4 Statistical analysis

Statistics were analyzed with SPSS statistics 22 (SPSS, Chicago, IL, USA), statistical difference was determined with an unpaired two-tailed analysis (T-test). Comparisons with *P* ≤ 0.05 were considered statistically significant. Results were expressed as means ± SD.

The pearson correlation coefficient between the fold change in RNA-Seq group and qRT-PCR group was determined by SPSS 22.0, one-way ANOVA followed by Duncan’s multiple range tests and differences were accepted as statistically significance when *P*<0.05.

## 3. Results

### 3.1 Body weight and liver histology changes

As shown in Table 1, compared with the control group, the bodyweight, food intake, specific growth rate (SGR) and feed conversion rate (FCR) of the experimental group rabbits were significantly decreased (*P*<0.05), while the hepatosomatic index (HSI) did not show significant differences between these two groups (*P*>0.05). During the experimental period, food intake of these two group did not show significant change (Fig. 2). The liver weight and the perirenal fat in the experimental group were significant decreased compared to the control group (*P*<0.05) (Fig. 3).

**Fig 2.**
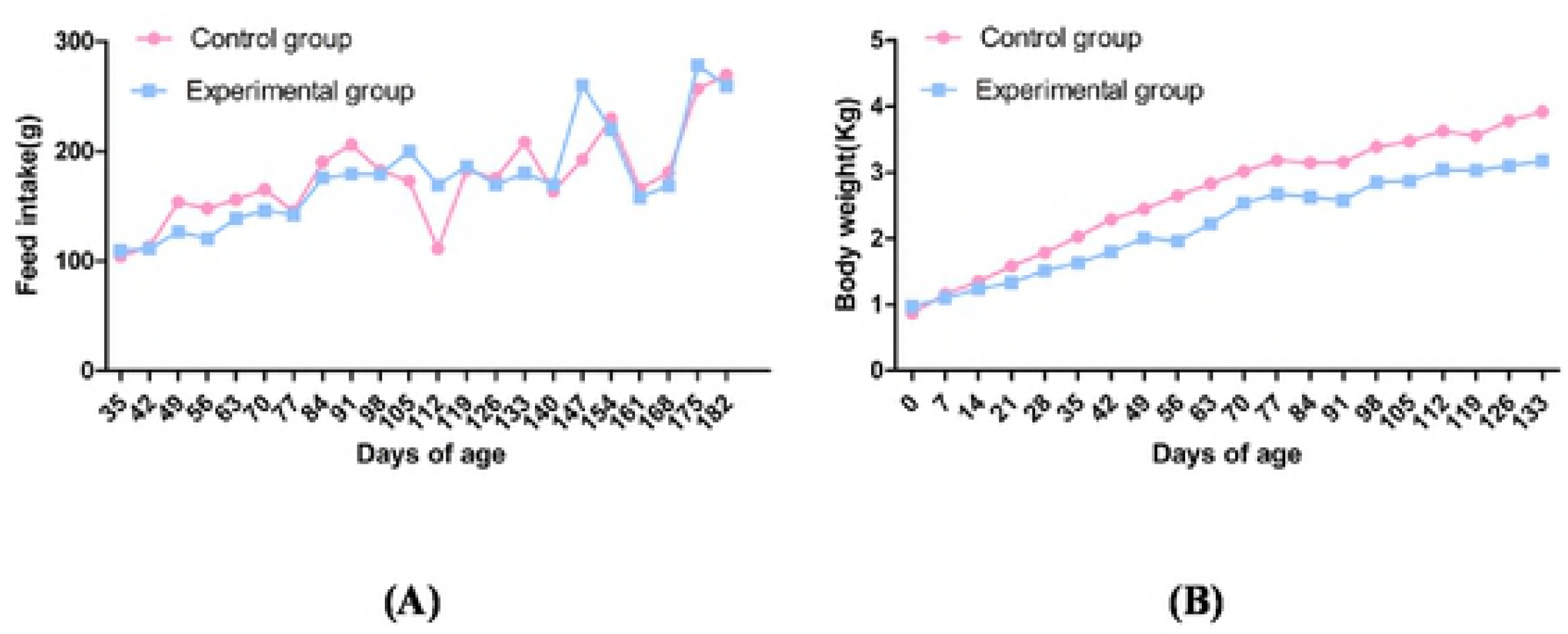
Changes of feed intake and body weight between the two groups. (A) Feed intake of the two groups. (B) Body weight of the two groups.

**Table 1.** Growth index changes of the experimental group and control group. In the same row, values with no letter or the same latter superscripts mean no significant difference (P>0.05), while with different small letter superscripts mean significant difference (P<0.05), and with different capital letter superscripts mean significant difference (P<0.01).

| Item | Treatment |  |
| --- | --- | --- |
|  | Control | Experimental |
| Initial weight (Kg) | 0.913±0.118<sup>a</sup> | 0.922±0.163<sup>a</sup> |
| Final weight (Kg) | 2.935±1.188<sup>a</sup> | 2.292±0.170<sup>b</sup> |
| Weight gain (%) | 2.021±0.159<sup>a</sup> | 1.370±0.262<sup>b</sup> |
| SGR (%day<sup>-1</sup>) | 0.033±0.002<sup>a</sup> | 0.022±0.004<sup>b</sup> |
| FCR | 3.555±0.341<sup>a</sup> | 4.795±1.238<sup>b</sup> |
| HIS (%) | 44.702±4.918<sup>a</sup> | 45.971±3.971<sup>a</sup> |

We also measured lipid droplet accumulation in liver tissue by using Oil red O staining, results showed that lipid droplet accumulation in the liver tissue of the experimental group is significantly low than the control group (*P*<0.05) (Fig. 4).

**Fig 3.** Weight change of internal organs between control group and experimental group. Experimental group and control group visceral weight (liver, perirenal fat, lung, kidney, heart and spleen) comparison in 90 d, Liver and perirenal fat differences were significant (*P*<0.05), lung, kidney, heart, spleen difference was not significant (*P*>0.05).

**Fig 4.** Changes of lipid droplet accumulation between two groups. (A) The liver section of control group. (B) The liver section of experimental group. (C) The quantified percentage of fat droplets in the two groups.

### 3.2 Serum biochemistry

As shown in Fig. 5, the HDL, LDL, TC, TG, TP and ALB did not show significant difference between the control group and experimental group(*P*>0.05).

**Fig 5.** Changes of blood biochemical parameters in the control and experimental group. (A) Serum biochemistry parameters of TC, HDL, LDL and TG. (B) Serum biochemistry parameters of ALB and TP. The blood biochemical parameters of the two groups. Results were analyzed by t-test.

### 3.3 Transcriptomic in the liver

A total of 14964 genes were expressed in liver tissues of 90-day-old New Zealand white rabbits using high-throughput transcriptome sequencing. In all, 444 DEGs were obtained between experimental group and control group, 262 of which were down-regulated and 182 were up-regulated in experimental group compared to control group (S2 Table). The DEGs had 63 GO terms (S3 Table) and were mapped onto 217 KEGG pathways (S4 Table). Among them, the most significant GO term was metabolic in the biological process and most of DEGs (5%) in retinol metabolism (Fig. 7B).

**Fig 6.** Characteristics of mRNA expression levels between the experimental group and the control group. Volcano map of differentially expressed gene.The point in DEGs volcano plot represent a gene, the abscissa indicates the logarithm of the difference in expression of the gene in two groups of samples, the larger the absolute value of the abscissa, the larger the difference multiple of the expression between the two samples; the ordinate indicates the negative logarithm of the statistical significance of the change in gene expression, The larger the ordinate value, the more significant the differentially expressed gene is. The green dots in the figure represent down-regulated DEGs, the red dots represent up-regulated DEGs, and the black dots represent indistinguishable genes. (B) Clustering map of differentially expressed genes. The abscissa represents the sample name and the clustering result of the sample, and the ordinate represents the clustering result of the differential gene and the gene. The different columns in the figure represent different samples, with different rows representing different genes. The color represents the level of expression of the gene in the sample log10. T01,T02,T03 were representative the control group,T04,T05,T06 were representative the experimental group.

**Fig 7.** The prediction and functional analysis of target genes regulated by differentially expressed mRNAs between the experimental group and the control group. (A) Differentially expressed genes KEGG classification map.The ordinate is the name of the KEGG metabolic pathway, and the ordinate is the ratio of the number of genes annotated to the pathway and the number of genes on the annotation. (B) Differentially expressed genes KEGG pathway enriched scatter plot. Each circle in the figure represents a KEGG pathway, the ordinate represents the pathway name, and the abscissa is the enrichment factor, which represents the ratio of the proportion of genes in the differential gene that are annotated to the pathway to the proportion of genes that all genes are annotated to. The larger the enrichment factor, the more significant the level of enrichment of the differentially expressed genes in this pathway. The color of the circle represents q-value, q-value is the *p* value after correction by multiple hypothesis test. The smaller the q-value, the more reliable the significance of the differential expression of the differentially expressed gene in the pathway; the size of the circle indicates the number of genes enriched in the pathway. Large circle indicate more genes.

**Fig 8.** qRT-PCR validation of 9 differentially generate from RNA-seq results in the liver of New Zealand white rabbit.

**Fig 9.** Pearson’s correlations between gene transcriptional level and phenotypic traits.

As shown in Figure 8, the mRNA expression of the selected genes including ABCA1, SREBP, CYP7A1, GPAM were identical to the results of the RNA-seq (Pearson’r=0.929). All the genes were related to the lipid metabolism (Fig.8).

In order to analyze the correlation between differentially expressed genes and phenotypic data, we obtained the numerical matrix through the heatmap visually by calculating their pearson correlation coefficients. The data information in the two-dimensional matrix is reflected by the color change. Among them, ACSM1, CYP7A1, SOAT2, APOA1 were positively correlated with TG, LDL and TP, and negatively correlated with perirenal fat, HDL, ALB and liver weight; ABCG5, SCD-1, ACSS2, CYP4A2, perirenal fat, HDL, ALB were positively correlation with liver weight, and negatively correlated with TG, LDL and TP (Fig. 9).

**Fig 10.** qRT-PCR validation of related genes in the retinol metabolism.

**Fig 11.** Pearson’s correlations between retinol metabolism gene and phenotypic traits.

## 4. Discussion

In the present study, growth performance results showed that banned the rabbit from eating their soft feces had great effects on body weight, weight gain, and SGR. After slaughter, weight of liver weight, tissue and perirenal fat weight of the experimental group was lower than the control group. Further analysis was performed to measure the lipid droplet accumulation in liver tissue between these two groups, Oil red O staining results of the liver tissue indicate that lipid droplet accumulation was decreased in the experimental group compared with the control group, so as the quantified data, Results in our present study is consistent with the previous study [35]. Some research showed that, under a normal diet condition, prevent the rabbits from eating soft feces leads them to develop malnutrition [36, 37]. It is speculated that there may be two main reasons for this phenomenon, one is that soft feces is rich in vitamins and microbial proteins, with the same feed intake, rabbits that were fasted to eating feces would consume 15% to 22% less protein. The other one reason is that soft feces of rabbits contain a large number of microorganisms, when forbid the rabbits from eating soft feces, resulting in changes in the microflora in their digestive tract and the reduction of bacterial populations [38, 39].

The soft feces eating rabbits can obtain 83% of niacin, 100% of riboflavin, 165% of pantothenic acid, and 42% of vitamin B12 from soft feces per day, which greatly improve the utilization of sodium and potassium. Study on rat showed that fasting soft feces can inhibit the growth of rats[40]. And R Franz found that the coprophagy was an important factor to explain the ringtail possum’s low requirement for nitrogen and its ability to subsist on a sole diet of Eucalyptus foliage[41]. Experiment showed that nutrient digestion rate of guinea pig’ is highly dependent on feeding of feces, soft feces that is eaten by the guinea pigs contain more nitrogen than the discarded feces[42]. The amount of methionine and lysine obtained from soft feces by adult beavers accounted for 26% and 19% of their total intake, respectively[43]. Altogether, these studies showed that nutrients in feces are critical for the growth and development of small mammals.

By using cloning and sequencing technology, Sánchez explored the main bacteria in the cecum of rabbits and found that 94% of the bacteria in the caecum are consisted with *Bacteroides*, there are a small amount of micro-bacteria, *Proteobacteria* [44]. By using CE-SSCP and qPCR techniques, Sylviecombes intended to determine the establishment of 2-day-old and 70-day-old rabbit cecal microbiota and found that the caecum flora evolved from a simple, unstable community to a complex, top-level community; The number of muramidomycetes reached to the maximum, and the total number of bacteria and the Bacteroides-Populus number reached to the highest values at 21 days of age[45]. The proportion of propionic acid decreased with age. The intestinal microflora and the host are mutually beneficial, and this pattern play an important role in physiological functions such as the host’s nutrient metabolism, immune system development, and disease resistance[46]. The importance of the intestinal flora has risen to a “super organ” which is necessary for humans and animals to survive and maintain their health[47].

Although much research have performed to study the effects of eating soft feces on the growth and metabolism of rabbits, their research are mainly focused on phenotypic data and serum biochemical indicators. Some of these studies indicated that the body weight of rabbits in the fasting feces group was lower than the normal group; the weight gain of the male rabbits was higher than the female rabbits between the two groups. There is no study aimed to investigate this phenomenon from the level of gene regulation. The effects of lipid metabolism on rabbits are expected to be explained at the molecular mechanism level. In the present study, the KEGG annotation indicated that different genes are mainly focused on retinol metabolism, pentose and glucuronide interactions, starch and sucrose metabolism, fatty acid degradation, steroid hormone biosynthesis. This showed that fasting feces has a certain effect on lipid metabolism in New Zealand white rabbits.

Liver is one of the most active organ for lipid metabolism, it is also a storage organ of retinol, more than 90% of the retinol is stored in the liver[48]. Retinol in the liver undergoes a biological role in the transport of plasma to peripheral target organs, and this process required the involvement of the specific vector retinol-binding protein (RBP)[49]. Studies have shown that RBP4 protein levels are positively correlated with TC and negatively correlated with HDL[50]. It is speculated that RBP4 enhances insulin resistance by regulating hepatic TC synthesis and VLDL release into blood, while RBP4 can affect fatty acid metabolism [51]. Disorders are closely related with insulin resistance. In our experiment, there was no difference in the expression level of RBP4 gene in the two groups, and the difference of TC and HDL were not significant, which imply that the prohibition of feces may not affect the expression of RBP4 gene in rabbit. Studies have shown that members of the cytochrome P450 family and ADH are involved in the process of retinol metabolism[52]. The CYP1 family catalyzes the conversion of retinol to retinal, a step that requires the participation of alcohol dehydrogenase (ADH) and ultimately conversion to retinoic acid[53]. Related studies have shown that knocking out of CYP1 family receptor down-regulated and the expression level of retinoic acid [54]. In this experiment, the CYP1A1, CYP1A2, and ADH4 genes in the experimental group were down-regulated compared with the control group, indicating that the prohibition of dietary feces may affect the metabolism of retinol. CYP26 is a key enzyme that regulates retinoic acid levels, degrading retinoic acid to an inactive product, and eventually excreted. Studies have shown that injection of CYP26A1 mRNA into Xenopus embryos and culture of embryos revealed symptoms of retinoic acid deficiency, suggesting that CYP26A1 is involved in the regulation of retinoic acid in vivo and the maintenance of normal embryo development[55]. Using the zebrafish embryo as a test material, the overexpression of Cyp26C1 showed the same symptoms as the overexpression of Cyp26A1 and the addition of retinal dehydrogenase inhibitors during embryonic period, indicating that Cyp26C1 is also involved in the process[56]. The metabolic process of folic acid. In this experiment, the CYP26 gene in the experimental group was down-regulated compared with the control group, indicating that the fecal behavior may lead to the loss of retinoic acid. Retinoic acid plays an irreplaceable role in embryonic development, organ formation, and proliferation and apoptosis of tissue cells[57]. Its lack or excess can cause abnormal organ development or embryo death, thus maintaining the balance of retinoic acid metabolism in the body. Importantly, it should not be regulated by diet alone. At the same time, attention should be paid to the activity of enzymes that synthesize and metabolize retinoic acid in the body, including RDH and RALDH of synthetic retinoic acid, and CYP26 activity closely related to retinoic acid catabolism.

Under the present condition, wearing a collar is a popular method to fast rabbit from eating soft feces, studies from other scholars indicated that wearing collars have no effects on food intake and drinking [58]. In the future, we plan to design a group of collars that allowed the rabbit to eat their soft feces to see its effects. Through studying the effects of fasting soft feces on New Zealand white rabbits, we can better understand the role of soft feces in the growth and development of New Zealand white rabbits, as well as to study the fecal behavior of other small mammals.

## 5. Conclusion

Fast eating soft feces reduced the body weight, liver histology and serum biochemistry of the New Zealand white rabbit compared to the control group. Through the RNA-seq technology, the results revealed with DEGs, GO and KEGG pathways. The metabolic disorder of retinol plays a key role in the growth and development of New Zealand white rabbits. While expanding the source of future studies with fat metabolism and meat quality traits, our findings provided more insights into mechanisms to enhance the productivity of animals.

## Acknowledgements

The authors are grateful to Meng Zhang provides guidance for drawing the figures, and Fang Li provides improvements in the test methods, Xiuling Li submitting the sequencing raw data online.

## Author contributions

**Conceptualization:** Ming Li and Guirong Sun

**Data curation:** Yadong Wang

**Formal analysis:** Tao Huang

**Funding acquisition:** Ming Li

**Methodology:** Guirong Sun and Jianhse Zhou

**Software:** Yadong Wang and Shuaijie Sun

**Supervision:** Ming Li

**Validation:** Yadong Wang and Mingming Xue

**Writing – original draft:** Yadong Wang

**Writing – review & editing:** Hui fen Xu and Juan J Loor

## Notes

The authors declare no competing financial interest.

## Data availability statement

All raw data have been deposited to NCBI Sequence Read Archive (SRA accession: PRJNA504548, Temporary Submission ID: SUB4763199).

